# Temporal refuges differ between anthropogenic and natural top down pressures in a subordinate carnivore

**DOI:** 10.1101/2020.06.09.143222

**Authors:** Rumaan Malhotra, Samantha Lima, Nyeema C. Harris

## Abstract

Apex predators structure communities through consumptive and non-consumptive pathways. In the carnivore guild, this can result in a within-guild cascade through the suppression of mesocarnivores. As the top-down influences of apex predators wane due to human-driven declines, landscape level anthropogenic pressures are rising. Human impacts can be analogous to apex predators in that humans can drive increased mortality in both prey species and carnivores, and impact communities through indirect fear effects and food subsidies. Here, we evaluate whether anthropogenic top-down pressures can structure communities in a similar manner as apex predators in shaping the interactions of mesocarnivores. Specifically, we expect anthropogenic forces to induce comparable effects as occurrence of apex predators in driving spatiotemporal partitioning between two mesocarnivores. Using multiple camera-trap surveys, we compared the temporal response of a small carnivore, the raccoon (*Procyon lotor)*, to the larger coyote (*Canis latrans)* at four sites across Michigan that represented opposing gradients of pressure from humans and apex predators. Contrary to our expectations, we found that raccoons shifted their activity pattern in response to coyotes at sites with higher anthropogenic pressures and exhibited no temporal response at sites with apex predators. Temporal shifts were characterized by raccoons being more diurnal in areas of high coyote activity. We conclude that despite superficial similarities, anthropogenic forces do not replace the function of native apex predators in structuring the mesocarnivore guild. As such, an intact and functioning native predator guild remains necessary to preserve spatiotemporal community structure, in natural and disturbed systems alike.

## INTRODUCTION

Predation impacts prey communities through consumptive effects, in the form of direct mortality (Sih et al. 1985, Matassa and Trussell 2011). However, predators can also structure communities through non-consumptive pathways, which can induce phenotypic plasticity in behavior (Lima 1998, Brown et al. 1999, Orrock et al. 2008, Prugh and Sivy 2020). The effects of non-consumptive pathways can be similar to those attributed to consumptive pathways, as numerous studies have shown analogous impacts (Lima 1998, Schmitz et al. 2004). For example, fear effects can be energetically costly, resulting in reduced fecundity, lower feeding rates, increased vigilance, and use of less profitable habitat to mitigate risks (Wirsing et al. 2008, Zanette et al. 2011). Similar to the effects of predation, varied resource use by prey induced by fear can initiate trophic cascades (Matassa and Trussell 2011). One of the best-known terrestrial cascades was driven in part by non-consumptive fear effects from an apex predator, the gray wolf (*Canis lupus*), in Yellowstone National Park, USA (Ripple and Beschta 2004). Fear effects are not exclusive to prey species; effects on smaller competitors are comparable (Durant 2000, Webster et al. 2012). Expanding the concept of mesopredator release, fear of large predators can also cause a trophic cascade by altering behavior of subordinate carnivores (Peckarsky et al. 2008, Suraci et al. 2016).

While fear effects are widespread in predator-prey interactions, they are not homogenous across the landscape. The landscape of fear is inherently based on the distribution of fear sources (i.e., higher-order predators) and habitat features that can buffer their threats (Oriol-Cotterill et al. 2015). For example, wild dogs (*Lycaon pictus*) respond to lion (*Panthera leo*) cues based on the density of vegetation, which relates to lion ambush potential (Webster et al. 2012). These non-consumptive effects are based also on predators’ behavior, and traits (i.e. speed, size) (Preisser et al. 2007, Cozzi et al. 2012). Changes in diet and space use by some species to avoid potentially deadly interactions suggests that fear effects are not experienced equally by subordinate species. Generalist species that are more plastic in certain aspects of their niche may be able to exploit refuges that minimize consequences of fear (Lima and Dill 1990, Forstmeier and Weiss 2004). Finally, multiple sources of fear may confound inferences of how subordinate species respond to a particular source of fear. Schuette et al. (2013) found that mesopredators responded to seasonal land use by humans, rather than inversely to occupancy by apex predators, contrary to expectations from mesopredator release hypothesis. While much research highlights variation in fear responses, it remains largely outstanding how discriminatory responses are to the particular source of fear or whether different sources can induce similar responses (Welch et al. 2017, Haswell et al. 2018, Lowrey et al. 2019).

Of particular interest in a rapidly expanding anthropocentric world is how human pressures compare to native fear sources such as apex predators. Cities are a rapidly growing, emergent habitat type with projected increases to 120 million ha globally by 2030 (McDonald et al. 2018). These human pressures increasingly drive the decline of apex predators at a global scale (Woodroffe 2000, Ripple et al. 2014). Similar to apex predators, humans can induce non-consumptive consequences on subordinate species through changes in space and time use (Ciuti et al. 2012, Clinchy et al. 2016). However, humans are unique in their top-down pressures in that they can exert fear effects across trophic levels, superseding hierarchies in natural systems (Smith et al. 2017, Suraci et al. 2019). The resultant heterogeneity of apex predator distribution from human pressures can induce differences in community structure as well as coexistence mechanisms within the carnivore guild (Berger 2007, Muhly et al. 2011). In urban areas, where spatial overlap among species are inevitable due to the limited amount of habitat available, temporal partitioning may be particularly important for species’ persistence (Adams and Thibault 2006, Santos et al. 2019). Therefore, given humans and apex predators can both exert fear effects, are responses comparable in subordinate mesocarnivores? Clinchy et al. (2016) provided some evidence that fear effects from humans exceeds that of native apex predators on badgers (*Meles meles*) in Britain (Clinchy et al. 2016). Additionally, Dorresteijn et al. (2015) reported that effects from human presence more strongly reverberate across trophic levels compared to the apex predators’ effects within a terrestrial ecosystem.

Urban-rural gradients provide natural experiments for comparisons of ecosystem function between natural and anthropogenic forces (McDonnell and Pickett 1990, Ellington and Gehrt 2019). Thus far, urban-rural gradients have predominantly highlighted changes in physical characteristics (e.g., body size) that can affect ecological interactions, or changes in biodiversity and species composition across taxa (Marzluff 2001, Urban et al. 2006). However, recasting the implications of urban expansion from the primary focus of degradation to evolutionary potential occurs by considering them as novel ecosystems that have conservation value (Kowarik 2011, Seto et al. 2011, Alberti 2015). We leverage and expand upon the urban-rural gradient formed by human pressure and the loss of apex carnivore presence to determine whether these different top-down forces induce similar spatiotemporal dynamics between a widely distributed subordinate carnivore and a larger sympatric competitor (Figure 1).

**Figure 1.**
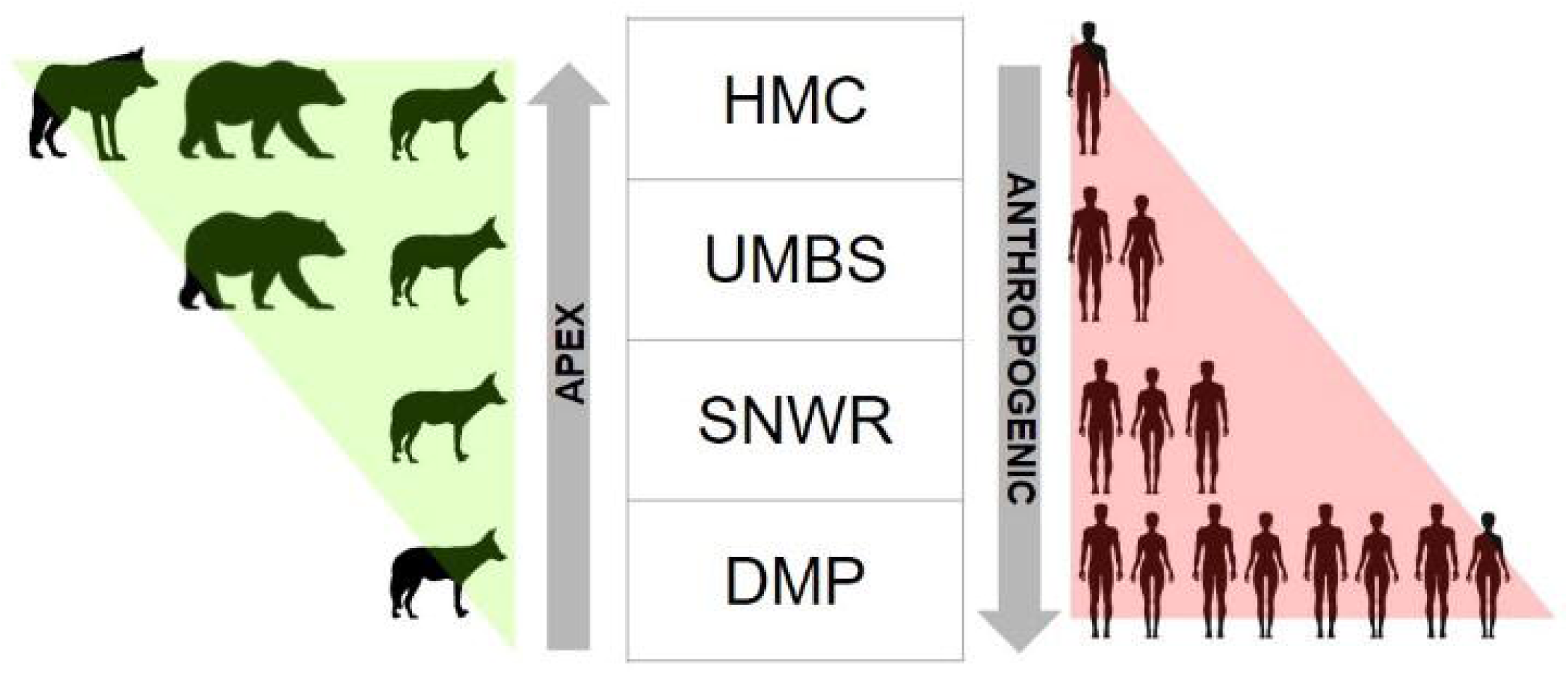
Opposing gradients in apex predation and anthropogenic pressures across sites, confirmed by photographic evidence during the study. From top to bottom, the Huron Mountain Club (HMC), the University of Michigan Biological Station (UMBS), the Shiawassee National Wildlife Refuge (SNWR), and the Detroit Metroparks (DMP).

Specifically, we test whether anthropogenic top down forces can impact the spatiotemporal interactions between a subordinate carnivore, the raccoon (*Procyon lotor*) and a large mesocarnivore, the coyote (*Canis latrans*) in a similar manner as natural top down forces induced by the presence of apex predators. Coyotes are known predators of raccoons and can have suppressive effects on raccoons in areas lacking gray wolves (Clark et al. 1989, Rogers and Caro 1998). We hypothesize that raccoons will shift their temporal activities in response to high activity zones of coyotes only at sites without gray wolves or heavy human pressure. In other words, top-down pressures, whether natural or anthropogenic, will suppress the dominance of coyotes on raccoons. Conversely, where these top down pressures are absent, raccoons will employ temporal refuges to reduce overlap with their dominant competitor. As anthropogenic pressures increase, our knowledge of contemporary baseline ecological interactions becomes dated. Thus, it becomes essential to understand how these competitive interactions compare across landscapes with varying human pressures.

## METHODS

### Study areas

We investigated raccoon temporal dynamics across differing levels of coyote activity at four sites across the state of Michigan, USA (Figure 2). The four sites encompass gradients of native (large predators) and novel (anthropogenic) top down forces (Figure 1).

**Figure 2.**
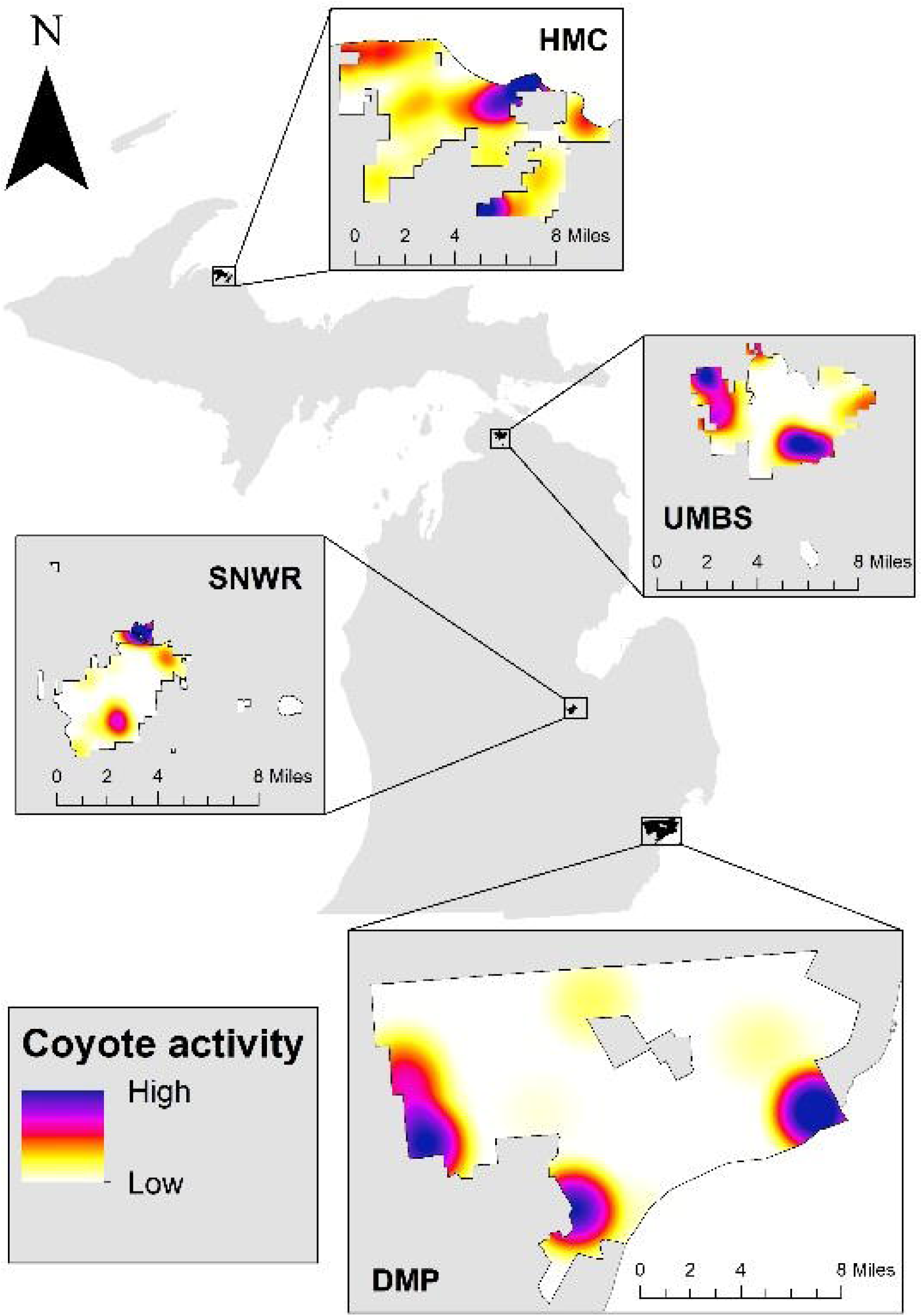
Kernel density heatmaps of coyote spatial use at the four study sites based on the number of independent coyote triggers at each camera. From north to south, the Huron Mountain Club (HMC), the University of Michigan Biological Station (UMBS), the Shiawassee National Wildlife Refuge (SNWR), and the Detroit Metroparks (DMP).

#### 1) Heavy apex pressure

The Huron Mountain Club (HMC) is a privately-owned property along the southern shore of Lake Superior, encompassing around 6,900 hectares in Marquette County, Michigan, USA. This site has a wide variety of habitats including beech-sugar maple hardwood forests, aspen dominated stands, and coniferous boreal forests. Sympatric large predators include: gray wolves, black bears (*Ursus americanus*), and coyotes. Anthropogenic pressures are limited to a small, seasonally occupied area of human habitation near the north central part of the property. Hunting and fishing occur on the property, and the intensity is presumably low due to restrictive public access.

#### 2) Relaxed apex pressure

The University of Michigan Biological Station (UMBS), a 4,000 hectare research station and forest in Pellston County, Michigan, USA served as one of our intermediate disturbance sites. With repeated logging and fire disturbance until 1923, the secondary forest is a mix of transitional hardwood and boreal forests. Douglas and Burt lakes along the north and south, and the town of Pellston and a major highway along west and east, respectively border this study area. Large co-occurring predators include: black bears, coyotes, and coyote-wolf hybrids. We were able to distinguish the few known coyote-wolf hybrids in the area due to them having collars from a different study, which were visible in the camera trap images (Wheeldon et al. 2012). Human pressures resulted from regulated research infrastructures for climate monitoring and housing facilities with low levels concentrated seasonally during the summer.

#### 3) Relaxed anthropogenic pressure

The Shiawassee National Wildlife Refuge (SNWR) is a 4,000 hectare wildlife refuge managed by the US Fish and Wildlife Service. The refuge is comprised of forested hardwood wetlands and lakeplain prairie. The city of Saginaw abuts the northern edge of the refuge and is surrounded by agricultural land for crop farming. The only large native predator present is the coyote. Anthropogenic pressures, in addition to the urban and ex-urban nature of the boundaries, are in the form of recreational visitors. Public hunting for deer and waterfowl, and furbearer trapping are permissible on the refuge in accordance with lawful seasons.

#### 4) Heavy anthropogenic pressure

The Detroit Metro Parks (DMP) is a collection of greenspaces interspersed throughout southeast Michigan that is managed by a city government department. We chose twenty-five of these parks that varied in size from ∼1.6-480 hectares, tree cover, human visitation, and degree of disturbance. Roads, buildings, or a riverine edge bound all parks. The only large native predator present is the coyote. Strong anthropogenic pressures are present in the form of the surrounding urban matrix, as well as the associated presence of humans and domestic pets across parks.

### Study species

Coyotes: As a highly successful mesocarnivore, coyotes exploit a wide range of habitats and exhibit tolerance to disturbance (Bekoff and Gese 2003, Flores-Morales et al. 2019). They exemplify mesopredator release through range expansion that aligns with human caused extirpation of wolves. Coyotes are subordinate to gray wolves where they are sympatric, and they are an aggressor species for several smaller carnivores, accounting for high rates of mortality for some species (e.g.,*Vulpes velox, Vulpes macrotis*) (Bekoff and Gese 2003, Berger 2007). As a result, coyotes are commonly cited as a species that can act as both a mesopredator or an apex predator in their community, depending on the presence of the gray wolf (Prugh et al. 2009, Roemer et al. 2009, Colborn et al. 2020). Although, Gehrt and Prange (2007) found no evidence of spatial avoidance by raccoons that would suggest avoidance of coyotes, they did not consider temporal partitioning as a strategy of avoidance (Prange et al. 2004).

Raccoons: Raccoons are dietary generalists that are also widespread across North America with much of their range overlapping that of the coyote (Timm et al. 2016, Kays 2018). They are found across a range of habitats from undisturbed forests to urban areas (Gehrt and Clark 2003). Raccons exhibit spatiotemporal variation in behavioral attributes, leading us to expect that the response of raccoons to coyotes may vary by differences in habitat and other characteristics across sites (Beasley et al. 2011).

### Camera trap survey

We deployed remotely-triggered camera traps (Reconyx© PC 850, 850C, 900, 900C) throughout each site with camera placement and sampling design proportional to study area size (Table 1). Our study uses data from one survey at DMP (2017-2018), two surveys at SNWR (2016, 2018), two surveys at UMBS (2015, 2016), and two surveys at HMC (2016, 2017-2018). We captured the heterogeneity of habitat and other environmental features to ensure ecological representation in the micro-site selection of camera traps. Camera traps were affixed to trees > 0.5m diameter and placed 0.5-1.0 m off the ground. Site-specific location of camera trap placement was determined by signs of animal activity such as game trails and scat. Camera trap settings included: high sensitivity, one-second lapse between three pictures in a trigger, and a 15-second quiet period between triggers. Camera traps were not baited.

**Table 1.** Temporal overlap (Δ) coefficients and 95% confidence interval for raccoon activity in low and high coyote zones within each camera survey in Michigan. Trap nights equals the total number of cameras multiplied by the number of nights each camera was active. NA indicates when survey lacked sufficient raccoon triggers to complete overlap analysis. Wald test for significance: ^+^ *p* < 0.1, * *p* < 0.05, ** *p*< 0.01.

| Survey period | Site/Year | Trap nights (n) | # Cameras | $\Delta$ (CI) |
| --- | --- | --- | --- | --- |
| Jan-May | HMC'18 | 3640 | 96 | NA |
| Nov-Mar | HMC'18 | 4200 | 96 | NA |
| Oct-Jan | HMC'17 | 2800 | 43 | 0.53 (0.21-0.82) |
| July-Dec | HMC'17 | 5739 | 43 | 0.67 (0.29-0.96) |
| Jul-Oct | HMC'16<sup>+</sup> | 8294 | 101 | 0.75 (0.59-0.89) |
| Sep-Dec | HMC'17 | 3885 | 43 | 0.76 (0.41-1.04) |
| Oct-Dec | UMBS'15 | 3320 | 47 | 0.82 (0.70-0.92) |
| Oct-Jan | UMBS'15 | 4447 | 47 | 0.83 (0.72-0.93) |
| Jul-Oct | UMBS '16 | 5612 | 57 | 0.86 (0.79-0.92) |
| Aug-Dec | UMBS'16<sup>+</sup> | 6275 | 57 | 0.86 (0.77-0.93) |
| Nov-Mar | DMP'18<sup>+</sup> | 4466 | 48 | 0.87 (0.59-0.89) |
| Jan-May | SNWR'16* | 5539 | 53 | 0.87 (0.84-0.91) |
| Jul-Dec | UMBS'16<sup>+</sup> | 8586 | 57 | 0.88 (0.83-0.93) |
| Sep-Dec | SNWR'18* | 3613 | 53 | 0.90 (0.85-0.94) |
| Oct-Dec | SNWR'18 | 3613 | 53 | 0.91 (0.85-0.97) |

Image identifications were initially crowd-sourced and filtered for carnivores using a public-science program called *Michigan ZoomIN* in combination with a consensus algorithm and expert validation (https://www.zooniverse.org/projects/michiganzoomin/michigan-zoomin). Carnivore species identifications were sorted and confirmed by at least two independent researchers in the Applied Wildlife Ecology Lab at the University of Michigan. These images and the associated metadata were used to assess raccoon diel activity and the spatial distribution of coyotes for analysis. Prior to all analyses, a 30-minute quiet period was introduced for every species to account for pseudoreplication, given the tendency of some animals to remain in front of the camera trap and trigger it multiple times.

### Analysis

We aim to determine if the interaction of the coyote and raccoon varied across sites in accordance with top-down pressures. Specifically, we test whether: a) raccoons were employing temporal partitioning in areas where they spatially overlapped with a high relative abundance of coyotes, and b) whether these results varied based on the higher order pressures present for the coyote (i.e. native apex predators, or humans). To do this, we had to:

1. Test for spatial partitioning at each site
2. Divide each site into areas with high and low relative abundance of coyotes
3. At each site, compare the temporal use of raccoons between areas with high and low relative abundance of coyotes
4. Compare the results of the temporal analyses between paired sites

### Testing for spatial partitioning

We used the Getis-Ord Gi* statistic (Z-score) in ArcGIS Pro to conduct pairwise tests of whether there was significant lack of overlap between coyotes and raccoons in each survey. Hot spots represent statistically significant (*pvalue* < 0.05) clusters of high numbers of species triggers (high relative abundance), while cold spots are statistically significant clusters with low numbers of species triggers (low relative abundance). If raccoons exhibit spatial avoidance, we expect that cameras designated as coyote hot spots would also be raccoon cold spots.

### Coyote relative abundance zones

We divided each survey into low and high coyote zone based on kernel density estimation from the number of independent coyote detections at each camera. Due to variation in camera spacing between sites, coyote activity zones were uniquely determined for each survey using quantiles of the survey data rather than a single cutoff value across all surveys. Cameras that fell within the bottom three quantiles of coyote triggers for each analysis period were assigned a low while those in the highest quantile were assigned to the high coyote zone. The bottom three quantiles were combined for the low designation to ensure adequate data for temporal analyses, as well as to specifically distinguish raccoon temporal dynamics in peak zones of coyote activity. Coyote triggers were checked for spatial independence using Moran’s I prior to kernel density estimation.

### Temporal analyses

We then used exact timestamps from raccoon detections to determine temporal distributions in low and high coyotes zones of activity. To evaluate if raccoon differentially use time with coyote presence, we calculated an overlap (Δ) coefficient of temporal activity for raccoons across coyote zones along with 95% confidence intervals generated from 10,000 parametric bootstraps of the temporal distribution models. We then used a randomization test with a Wald statistic to determine the probability that the two sets of circular observations from zones arose from the same distribution (Ridout and Linkie 2009, Sovie et al. 2019). Δ values range from 0 to 1, with 0 indicating completely distinct and non-overlapping temporal activity between comparison groups, and 1 indicating complete overlap. Δ_1_ was used for comparisons when one of the sample groups had less than 50 triggers; otherwise Δ_4_ was used to estimate temporal overlap (Ridout and Linkie 2009). Activity patterns pairs for randomization tests that yielded a significance or near significance (*pvalue* < 0.1) were also visually assessed to determine how the peak activity shifted (i.e. primarily nocturnal to crepuscular). Furthermore, all analysis results which fell within significance thresholds were further tested for overlap between coyote and raccoon activity within each high coyote zone using the same methodology (generating overlap coefficients, and comparing using a Wald test). This was to determine whether raccoons were exhibiting temporal shifts due to temporal avoidance of coyotes. All temporal analyses were conducted using the ‘overlap’ and ‘activity’ package in program R. Spatial analyses including kernel density estimates, Getis-Ord Gi* Z-scores and Moran’s I were implemented in ArcGIS Pro (vers. 2.3.1).

### Accounting for seasonal and yearly variation

Our multi-site camera study allowed us to compare differences in raccoon response to coyote based on landscape level differences in native apex predator and anthropogenic pressures. First, we determined overlap in raccoon activity between low and high zones of coyote presence for each survey. This gave us a measure of within-site variation in raccoon temporal activity. Next, we used these within-site measures of raccoon temporal activity to conduct pair-wise comparisons with the other sites, which resulted in a single survey being used in multiple different pairwise analyses (Figure S1). For example, the UMBS 2016 survey was compared to the HMC 2017 survey as well as to SNWR 2018. Lastly, we aggregated the results of all the pairwise analyses between sites to examine recurring patterns (i.e. were there significant shifts in time use by raccoons along an anthropogenic gradient or at the extreme opposites?). Comparing between seasons can confound inferences from the analyses, due to different seasons potentially resulting in different detection rates (Rowcliffe et al. 2011). Thus, to account for differences in seasonality, we only used data from the overlapping time period for every pairwise site comparison (Figure S1). For example, we subsetted the data from SNWR and UMBS to mid-October through mid-December to compare raccoon diel shifts across coyote zones, as this is the overlapping period even though the individual surveys were longer. We also compared the inter-annual variation in activity patterns of both raccoons and coyotes at UMBS and HMC to ensure that there was not major variation in results between years. Note: this was not possible for SNWR and DMP, as we did not have replicate surveys in the same season at those sites that occurred between different years.

## RESULTS

We obtained 1,079 coyote and 7,502 raccoon triggers with a 30-minute quiet period from 2015-2018 across eight surveys in 58,459 trap nights (HMC-31,808; UMBS-13,033; SNWR-9,152; DMP-4,466). Raccoons and coyotes were the most common carnivores in almost every survey, comprising 57-98% of all the carnivore triggers. In Detroit, where domestic dogs and cats comprised 35% of the triggers, coyotes were the fourth most common carnivore species after raccoons, cats, and dogs. To account for the variation in survey season, we used only seasonally overlapping surveys for analyses (Figure S1). Subsets of each survey that overlapped completely in days of the year were used to make between site comparisons (mean = 100 days; range = 53 – 146 days). Expected top down natural pressures for each site were confirmed via camera trap images with both wolves and black bears occurring only at HMC, and black bears and coyote-wolf hybrids at UMBS. Likewise, more anthropogenic pressures in the form of human presence occurred at SNWR and DMP, with coyotes as the largest predator at both these sites. Low numbers of triggers for wolves, coyote-wolf hybrids, and bears prevented analyses exploring how these apex predators influence raccoon activity.

### Spatial partitioning

Getis-Ord Gi* statistics indicated that coyotes were distributed non-randomly in space. Hot spots and cold spots of both species were not consistent within sites; statistically significant clustering (*pvalue* < 0.05) varied at each site by survey. Hot and cold spots for both species were present in some surveys and absent in others. In other cases, hot spots shifted between surveys and years to opposite ends of the site. Of the 28 coyotes cold spots identified across surveys, none overlapped with raccoon hot spots. Moreover, the 13 raccoon cold spots did not coincide with coyote hot spots for a single survey, indicating that raccoons were not spatially avoiding areas of high coyote abundance (Table S1). We even applied a more lenient threshold for significance (*pvalue* < 0.1) and results did not change.

### Coyote relative abundance

Kernel density estimates corroborated Getis-Ord Gi* statistic results that indicated coyotes were distributed non-randomly in space. At DMP with heavy anthropogenic pressure (average 77 coyote triggers per camera in “HIGH” coyote zones), coyote activity was concentrated in two heavily forested parks and had few human triggers compared to the rest of the surveyed parks in Detroit. In contrast, at HMC with heavy natural apex pressure, highest coyote activity occurred in a recreation area that contains several buildings and homes but had fewer overall triggers (average 3 coyote triggers per camera in “HIGH” coyote zones). Coyote activity formed distinct zones in the intermediate sites as well, although these hotspots were not associated with any discernible landscape level measures of anthropogenic pressures.

### Site dynamics (Table 1)

Raccoon triggers were recorded within both the low and high zones of coyote activity across all sites, establishing spatial overlap between the two species (Figure S2). Again, we expected that top-down pressures, whether natural or anthropogenic, would suppress the dominance of coyotes on raccoons resulting in no temporal shift at HMC and DMP.

#### 1) Heavy apex pressure (HMC)

In total, we recorded 517 and 75 raccoon triggers in the low and high coyote zones, respectively. Neither survey considered for HMC in 2017 or 2018 showed evidence of raccoons shifting temporal use, as expected. However, low number of raccoon triggers in high coyote zones limited our assessment of temporal shifts. In contrast, the HMC 2016 survey had more triggers (N_LOW_ = 209, N_HIGH_ = 28) and showed a moderate shift in raccoon temporal activity between zones (Δ_1_ = 0.79, CI: 0.586-0.892, *p* = 0.081). Raccoon activity was bimodal during night-time in low coyote zones, exhibiting a distinctive larger peak around the beginning of the nocturnal period. In heavy coyote activity zone, raccoon temporal activity peaked distinctly around the early morning.

#### 2) Relaxed top down pressure (UMBS)

We recorded 667 and 329 raccoon triggers in the low and high coyote zones, respectively. Contrary to expectations, for the majority of the survey periods considered, raccoons did not shift their temporal use in response to heavy coyote presence at UMBS. The only exception occurred for UMBS 2016 survey across the longest comparison period (146 days) with raccoons shifting temporal use (Δ_1_ = 0.88, in zones of heavy coyote activity CI: 0.827-0.926, *p* = 0.055). In comparison to the raccoon activity in the low coyote zone, which was nocturnal with a defined peak around the beginning of the nocturnal period, raccoons in the high coyote zone shifted peak activity to closer to midnight, and also exhibited increased crepuscular activity (Figure 5).

**Figure 3.**
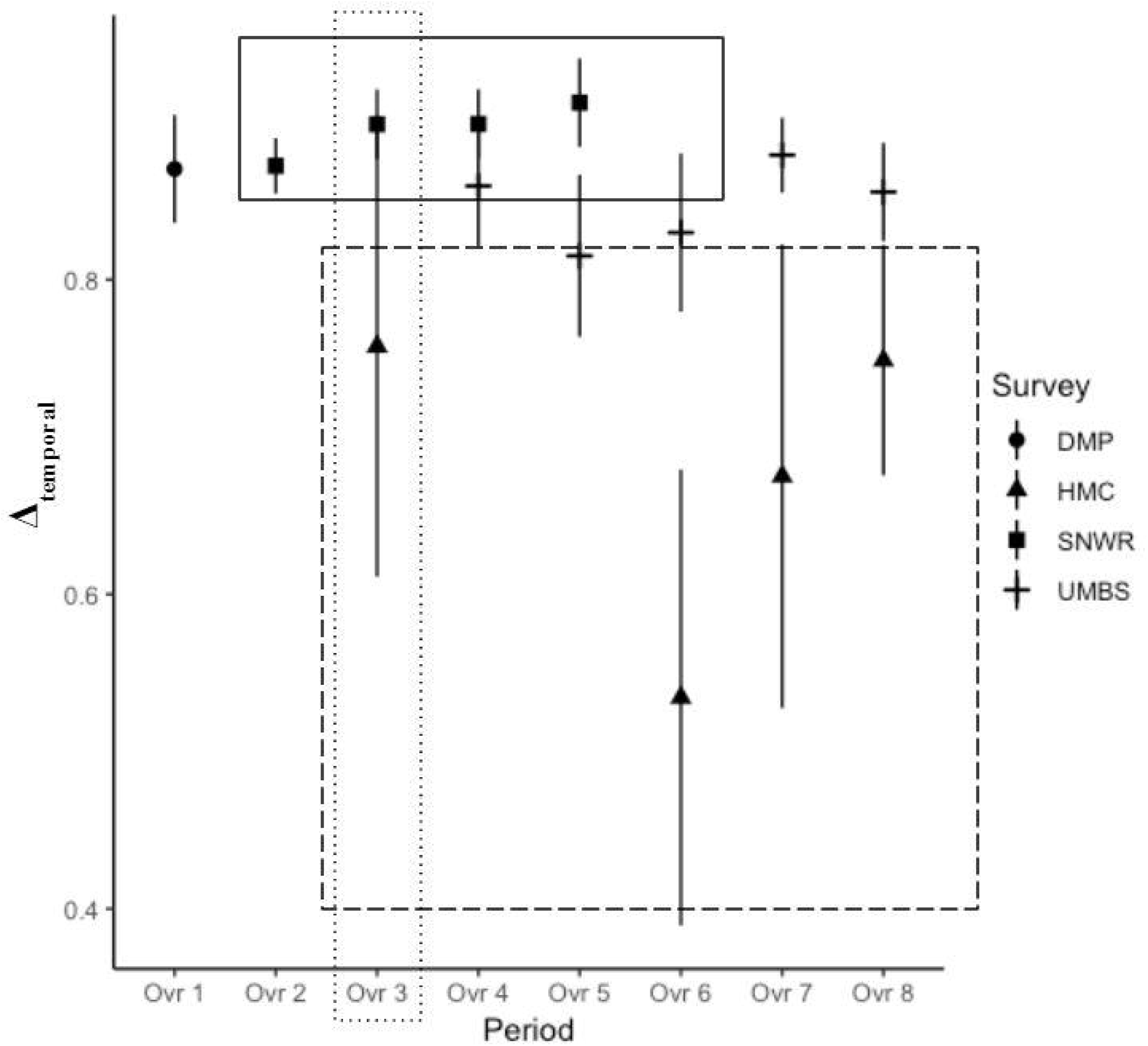
Mean temporal overlap (Δ_temporal_) in raccoon activity between high and low spatial zones of coyote activity with one standard error. The solid box represents sites with higher anthropogenic pressures, while the dashed box represents sites with higher apex predator pressures. The dotted box highlights one of the eight paired between-site comparisons.

**Figure 4.**
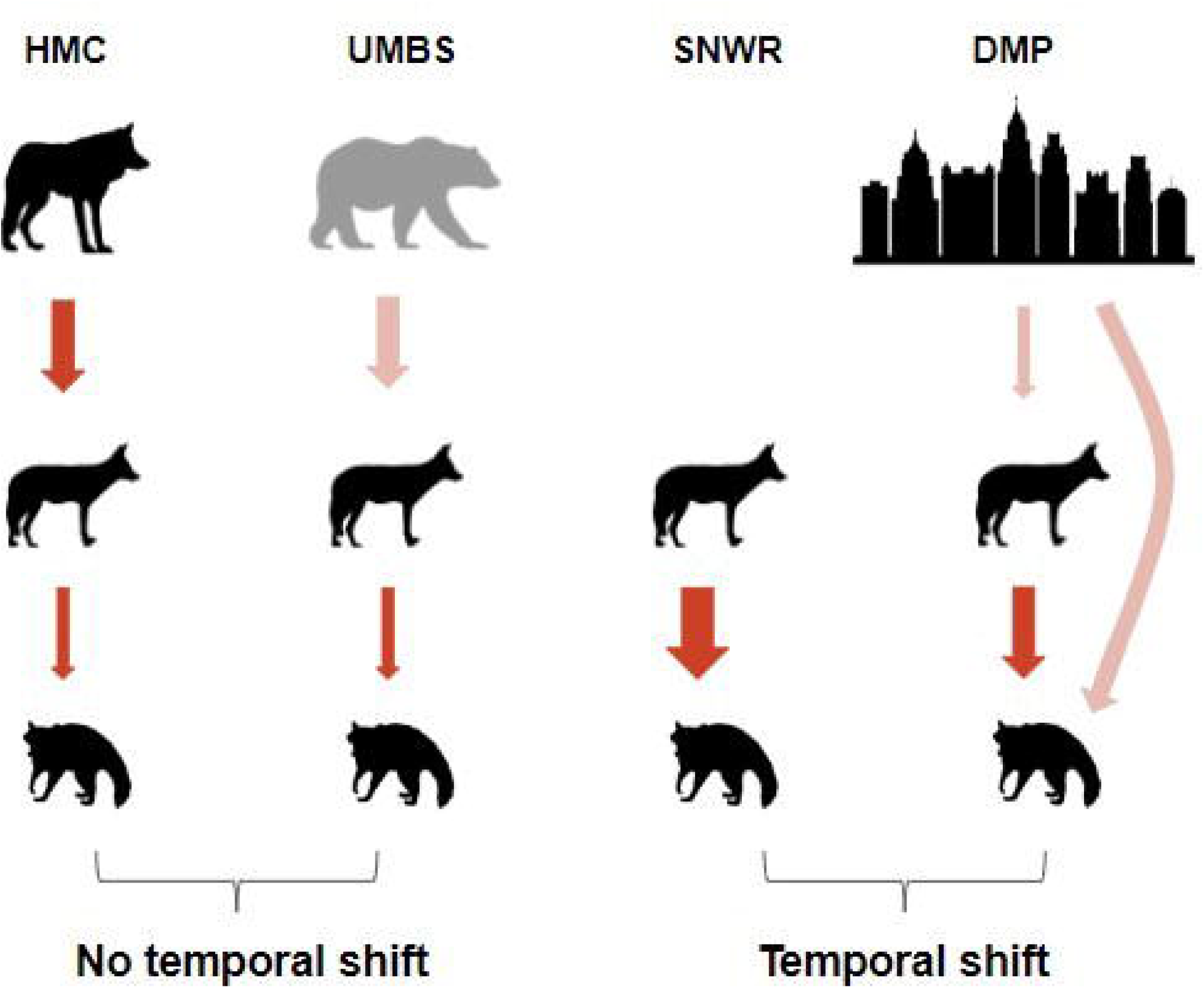
Spatial variation in diel shifts associated with coyote zones across Michigan study areas. Arrows represent hypothesized top down effects through the guild. Faded arrows represent hypothesized explanations specific to the results of this study. Star represents sites where temporal shifts was expected from top down pressures.

**Figure 5.**
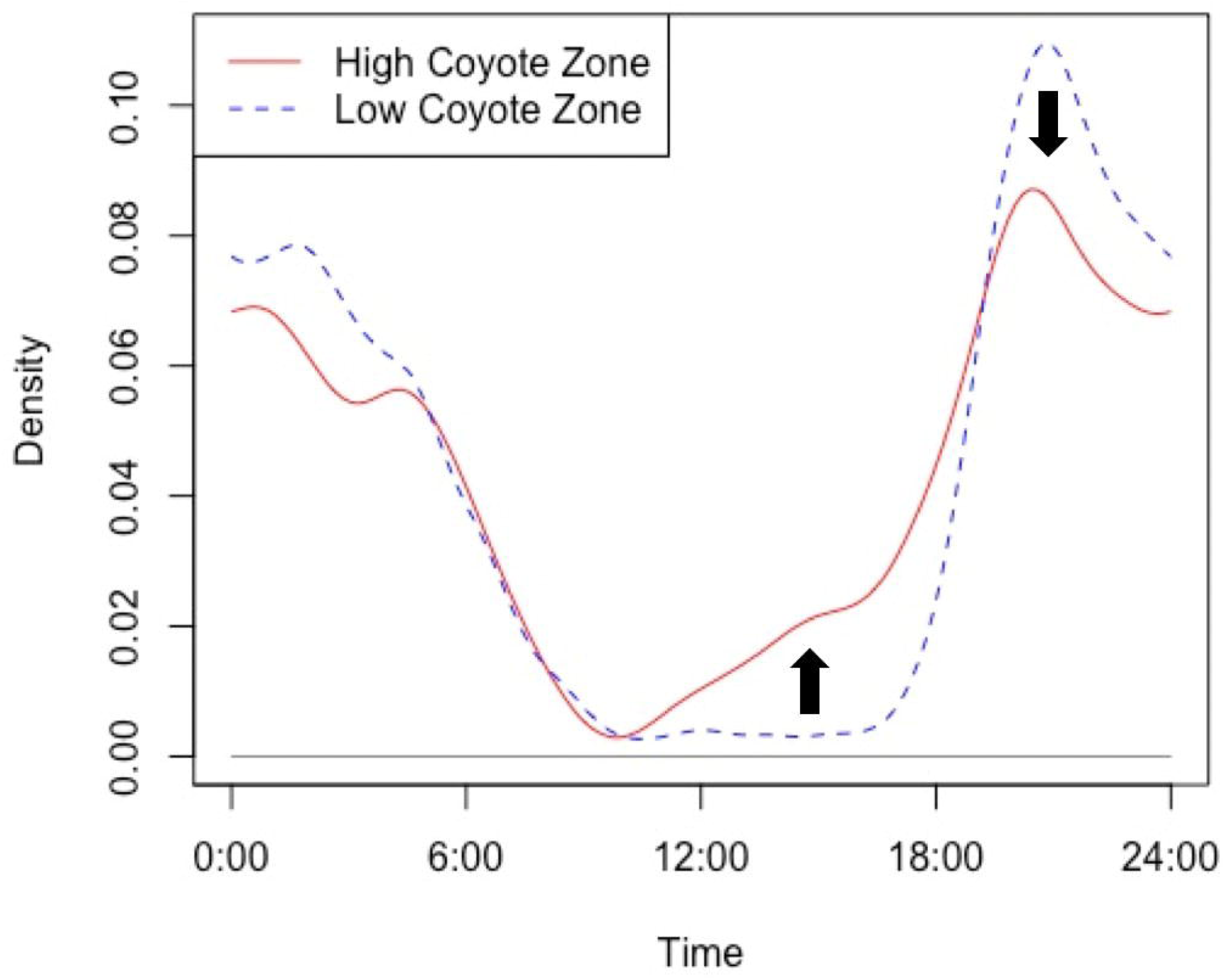
Characteristic shift in raccoon temporal activity in zones of high coyote pressure. The arrows represent characteristic shifts in every comparison where there was a significant or near significant difference in raccoon activity (p < 0.01), namely a loss in the distinctive peak at the beginning of the nocturnal period, and a marked increase in diurnal activity.

#### 3) Relaxed anthropogenic pressure (SNWR)

We recorded the highest raccoon activity at SNWR with 5377 and 1335 triggers in the low and high coyote zones, respectively. Raccoon activity in low coyote zones was similar to that of UMBS, being primarily nocturnal with a defined peak near the beginning of the nocturnal period. Similarly, in high coyote zones, the peak activity shifted closer to midnight, with increased crepuscular activity. However, in contrast with UMBS, nearly all the survey periods considered in SNWR showed significant shifts in raccoon activity in high coyote zones. We detected significant shifts in temporal activity for raccoons in three of the four comparison periods for SNWR surveys. This finding supports our expectations that raccoons would require temporal refugia at a site where coyotes are released from strong top down pressures. The only survey where raccoons did not shift their diel activity with coyote activity occurred in 2018 fall/winter (Δ_4_ = 0.91, CI: 0.846-0.9678, *p* = 0.538).

#### 4) Heavy anthropogenic pressure (DMP)

We recorded 282 and 93 raccoon triggers in the low and high coyote zones, respectively. Contrary to expectations, we found that raccoons adjusted time use (Δ_4_ = 0.87, CI: 0.795-0.938, *p* = 0.051) in heavy coyote presence. With heavy coyote presence, raccoon activity was more dispersed throughout the night, and included activity in the late morning. In low coyote density zones, raccoon activity showed a sharp peak around the beginning of the nocturnal period but was otherwise confined to the nocturnal period.

Generally, raccoons at sites with greater anthropogenic pressures (DMP, SNWR) exhibited shifts in temporal activity between low and high coyote zones. In contrast, raccoon and coyote exhibited consistent spatiotemporal overlap at sites with higher apex predation (HMC, UMBS), reflecting a rural-urban gradient. Furthermore, mean Δ overlap in raccoon low and high zones of coyote activity was higher with anthropogenic pressure sites compared to high apex predation sites (Figure 3). All shifts in raccoon activity from low to high coyote zones across sites that approached significance (*p* < 0.1, see Table 1) were characterized by the loss of strictly nocturnal activity patterns (Figure 5). Specifically, we observed the absence of a distinctive peak of activity at the beginning of the nocturnal period (19:00 hours).

For the results that approached significance (p < 0.1 or less), we tested whether shifts in raccoon behavior were due to reduced overlap with coyotes, using the same methodology used to compare raccoon activity between coyote zones (a Wald test). With the exception of HMC, we found that in high relative abundance coyote zones, raccoon temporal activity was significantly different from that of coyotes in every case (Table 2). This indicates that raccoons have reduced temporal overlap with coyotes, and is evidence for temporal partitioning.

**Table 2.** Temporal overlap (Δ) coefficients and Wald test results comparing coyote and raccoon activity in coyote high abundance zones for surveys where raccoon activity significantly (*p* < 0.1) differed between coyote high and low zones.

| Overlap period | Site | $\Delta$ | *pvalue* |
| --- | --- | --- | --- |
| Ovr1 | DMP | 0.78(0.69-0.86) | **0.0029** |
| Ovr2 | SNWR | 0.70(0.63-0.78) | **0.0001** |
| Ovr3 | SNWR | 0.73(0.64-0.82) | **0.0000** |
| Ovr4 | SNWR | 0.73(0.64-0.82) | **0.0000** |
| Ovr4 | UMBS | 0.74(0.67-0.85) | **0.0104** |
| Ovr7 | UMBS | 0.76(0.68-0.83) | **0.0000** |
| Ovr8 | HMC | 0.78(0.67-0.89) | 0.3150 |

### Yearly and seasonal variation

Prior to comparing results between sites, we tested for interannual variation in raccoon temporal use in response to coyote activity within sites. Two of our sites were surveyed in two different years with overlapping seasonality: UMBS (2015 and 2016), and HMC (2017 and 2018). Both sites showed similar results between years; no evidence of a shift in raccoon activity between high and low coyote activity zones.

To make inferences about the differing levels of apex and anthropogenic pressures inherent to each site, we had to compare raccoon temporal response to coyotes between sites during overlapping time periods (Figure 3, S2). For HMC-UMBS site comparisons, we found no evidence of a temporal shift at either site during the fall/winter period while weak evidence of a temporal shift at both sites during summer/fall period (HMC:Δ_4_ = 0.75, CI: 0.586-0.892, *p* = 0.087; UMBS: Δ_4_ = 0.88, CI: 0.827-0.926, *p* = 0.055). Raccoons at SNWR generally exhibited a significant shift in almost every comparison period, while comparable surveys at UMBS and HMC showed no shift in raccoon temporal activity. The DMP survey overlapped with and was compared to the survey at HMC. While DMP showed some evidence of a temporal shift (Δ_4_ = 0.87, CI: 0.795-0.938, *p* = 0.051), the low number of coyote triggers for that HMC survey prevented a comparison between the sites.

## DISCUSSION

Behavioral adjustments that reduce competition for resources prove necessary to promote coexistence (Inouye 1978). These shifts can be in diet, spatial or temporal use. We measured the use of temporal refuges by raccoons in the presence of coyotes across opposing sources of top down pressures, including gradients in the presence of humans and apex predators. As expected, we found that temporal shifts can occur when there is spatial overlap between the competitor species. More importantly, we highlight these temporal refuges vary with the top down pressures present at a site level. Generally, we found that at sites with higher anthropogenic pressures (SNWR, DMP), raccoons exhibited behavioral plasticity by shifting their temporal activity in response to coyotes. These results complement recent findings that: a) non-consumptive effects impact the spatial use within the carnivore guild (Newsome and Ripple 2015); and b) that non-consumptive effects (fear effects) are present within the hierarchy of the carnivore guild (Gordon et al. 2015).

We expanded our understanding of these phenomena by exploring whether a plastic response in the temporal niche of a subordinate carnivore (raccoon) was dependent on the source of fear (anthropogenic versus apex predator) exerting on the larger mesocarnivore (coyote). We found that while non-consumptive fear effects can cascade within a guild, anthropogenic and apex predator fear sources did not elicit a similar response from raccoons in areas of high relative abundance of coyotes. Instead, a temporal response by raccoons to high coyote presence followed an urban-rural gradient, such that raccoons shifted their temporal use in zones of high coyote presence in areas with more anthropogenic pressure (i.e., in or in close proximity to urban environments). This implies that humans may exert only weak top-down pressures on coyotes, resulting in raccoons still needing to be responsive to their dominance. Contrary to this finding, we expected no temporal shifts in raccoon activity at either site with top down pressures (HMC with apex predators or DMP with humans) due to the compromised function of coyotes.

Unlike HMC, the UMBS site does not have wolves. However, other large predators including black bears and coyote-wolf hybrids were captured on camera during surveys and could act as fear sources for coyotes. Therefore, the presence of apex predators at both HMC and UMBS indicates that raccoons are exhibiting temporal partitioning only where coyotes have ascended in trophic level at SNWR and DMP (Colborn et al. 2020). Similarly, Wang et al. (2015) found that raccoon detections decreased with coyote activity, but only at sites where pumas were absent, supporting mesopredator release (Wang et al. 2015). Even within zones of low coyote activity, raccoons are unable to avoid coyotes spatially due to the smaller patch sizes and ‘harder’ bounds of city parks (i.e. green space “islands” surrounded by roads and buildings). Furthermore, Breck et al. (2019) found that coyotes at urban sites are bolder in comparison to their rural counterparts, which would further support their role as a fear source at our DMP site. When viewed with the consistent shift in raccoon activity across SNWR with low anthropogenic pressure, we begin to conclude more generally that sites with higher anthropogenic pressures are also sites where coyotes exert stronger pressures on subordinate carnivores.

Fear effects are not static and can change with shifts in climatic conditions that define seasonal changes (Riginos 2015). Resource availability, such as the abundance of small mammal prey, fluctuates with season and could be a driver of varying levels of competition between coyotes and raccoons (Batzli 1992, Fedriani et al. 2000, Neale and Sacks 2001). At an urban site (e.g., DMP), food subsidies in the form of trash could reduce seasonal variation in resource competition (Oro et al. 2013, Newsome and Ripple 2015). The presence of these additional food sources could potentially offset the seasonal variation in natural sources of food (Oro et al. 2013). Thus, we would expect patterns of temporal use, particularly in the presence of a competitor, to vary seasonally (Sovie et al. 2019). Indeed, temporal shifting appeared seasonally dependent, with raccoons exhibiting changes in diel activity within coyote high zones during every survey period that occurred during the spring and summer months. Pairing dietary studies that explore the seasonal variation in coyote and raccoon diets across all sites with spatiotemporal analyses would elucidate if seasonal variation in resource availability drives resource partitioning between these species. In addition to factors affecting the temporal interactions of species, previous studies have shown that winter surveys can yield lower detection rates for species (Gompper et al. 2006, Ikeda et al. 2016).

Given that we did not quantify the amount of top down pressure from apex predators and humans or measure resource abundance at sites, we cannot discount the possibility that factors other than top down forces drove the urban-rural gradient we see in our results. Sites varied in vegetative cover, topography, latitude, and distribution of resources. However, differences in the sources of top down forces are the most obvious and likely ecological factor that differs between the sites for generalist species such as raccoons and coyotes. Our results highlight broad patterns in raccoon temporal use between zones of high and low coyote activity. The mechanisms that underlie these patterns require further study, and a temporal shift could very likely have more nuance than simple avoidance by a subordinate carnivore. For example, a shift in temporal use by a subordinate (as shown in our SNWR and DMP sites) might reflect tracking of an alternate resource to indirectly avoid competition with a larger competitor rather than direct avoidance of antagonistic interactions. This shift would not necessarily manifest as reduced temporal overlap between the competing species, and would require further studies quantifying the abundance of different resources. While our results indicate the response of the raccoon to be driven by a larger predator, it does not preclude an interaction between top-down and bottom-up forces, which may be important to understanding what raccoons are directly responding to (Elmhagen and Rushton 2007). Thus, we define “suppression” broadly as one species driving a shift in niche use of another species, without inferring the finer mechanisms operating across sites.

In the broader community context, we conclude that apex predators suppress the ecological function of coyotes to a greater degree than anthropogenic forces. The strong non-consumptive effects that humans exert may drive all members of the guild to avoid times of peak human activity, regardless of size hierarchies (Wang et al. 2015). Despite the presumed top down effects that humans can exert (Dorresteijn et al. 2015, Clinchy et al. 2016, Suraci et al. 2019), they do not inherently replicate the suppressive role that apex predators play for mesocarnivores, and the subsequent indirect effects for smaller carnivores. Areas with high anthropogenic pressures have suppressive effects across the guild, and yet the suppressive role of the coyote on a subordinate carnivore was maintained even in these heavily disturbed landscapes. The paradigm in conservation is shifting to include *in situ* conservation of species in urban habitats, rather than considering these areas solely as suboptimal sink habitats (Magle et al. 2012, Athreya et al. 2013, Mormile and Hill 2017). Studies comparing the ecological roles of species within a community between urban and natural systems are timely. Such work will prove invaluable in understanding how wildlife communities in these novel habitats differ not just in composition, but also in their function.

## ACKNOWLEDGMENTS

First, we recognize implementing our camera survey and field research was conducted on lands originally belonging to the People of the Three Fires. This work was made possible by the generous support and permission of the Detroit Metro Parks, the Shiawassee National Wildlife Refuge, the University of Michigan Biological Station, and the Huron Mountain Club. We would like to thank the Applied Wildlife Ecology Lab at the University of Michigan for assistance with fieldwork, image classification and overall project feedback. In particular, K. Mills and S. Gámez provided valuable commentary and general edits. We also thank the countless volunteers for assistance with fieldwork and Michigan ZoomIN. Funding was provided by the Detroit Zoo, the Huron Mountain Wildlife Foundation, and Ecology and Evolutionary Biology Department at the University of Michigan.

