## Supplementary material for "Temporal refuges differ between anthropogenic and natural top down pressures in a subordinate carnivore": Table S1

**Figure S1.** Concurrent surveys between sites used in analysis. Eight overlap periods (Ovr#) were determined by start and end dates that were sampled during both surveys to maximize the overlap period. The area between the dashed lines for each overlap represents the data used for spatial and temporal analyses to compare the two sites represented.

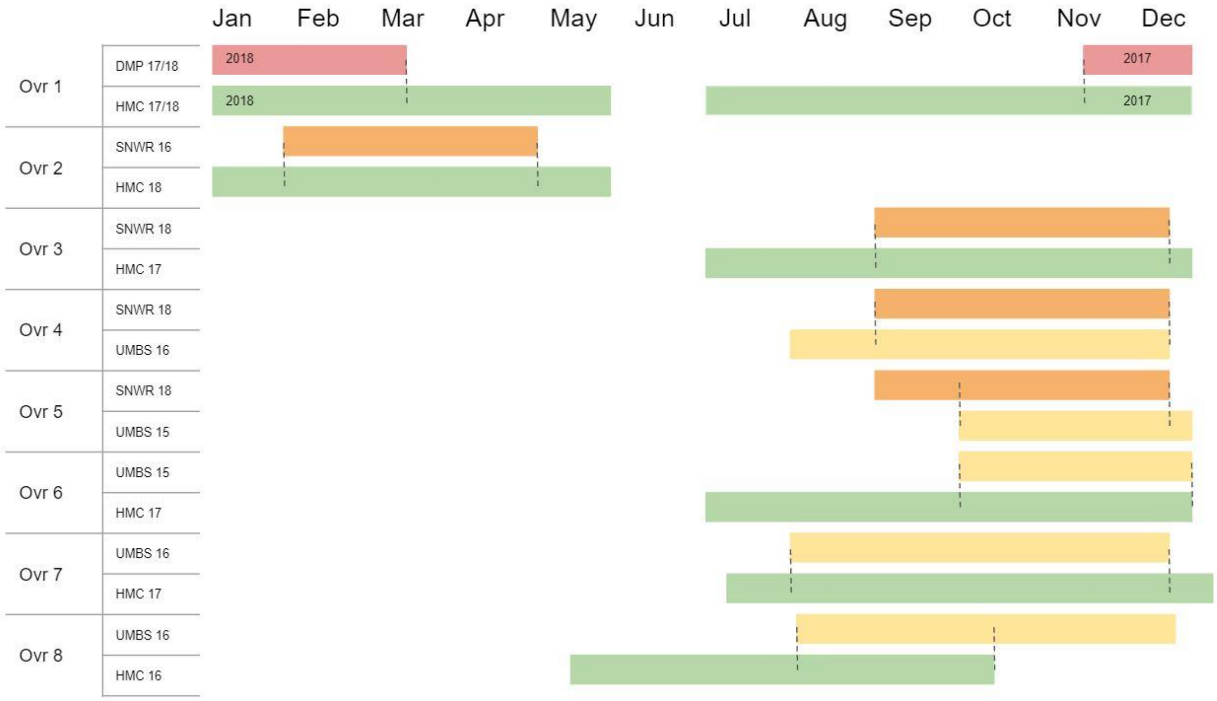

**Figure S2.** The proportion of raccoon triggers at each site across the two spatial zones of coyote activity.

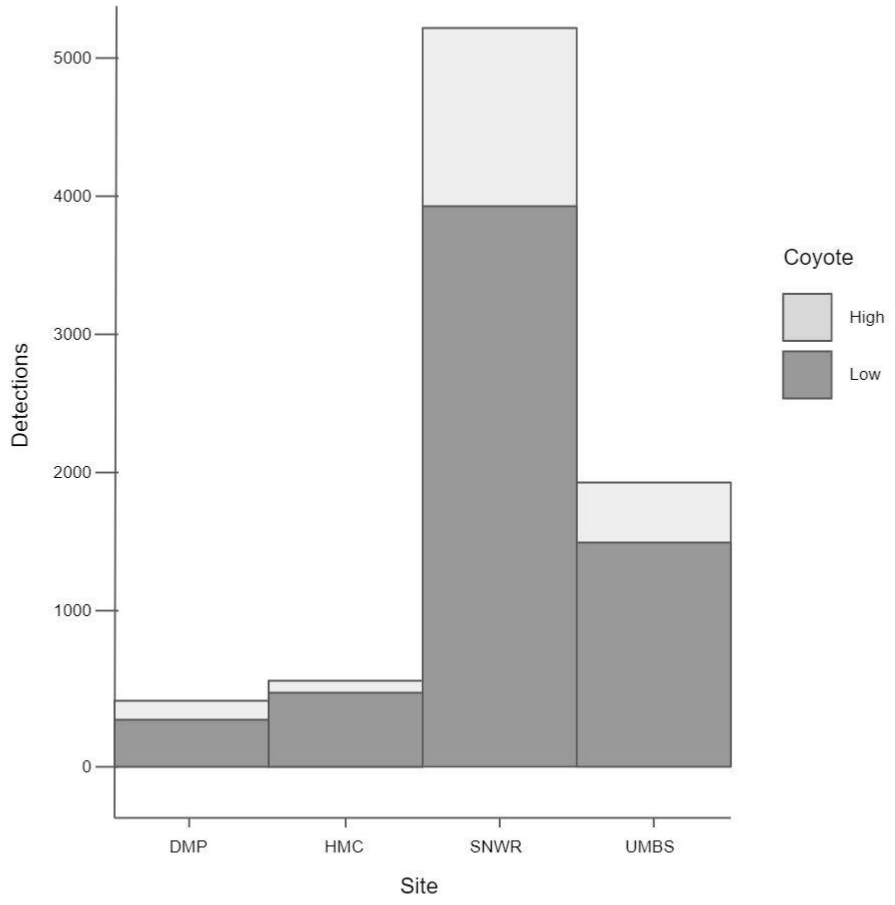

Table S1 Every significant hot or cold spot for each species, obtained through Getis-Ord Gi* statistic analysis. Each row represents a single camera during a survey of a specific site (designated under the ‘Site’ header). Hot spots were designated as Z-scores >1 and *pvalue < 0.1*. Cold spots were designated as Z-scores <1 and *pvalue < 0.1*. The coyote relative abundance column refers to the spatial zone designation for each camera, obtained kernel density estimation. Colors are scaled based on significance ranging from *pvalue* < 0.1 to *pvalue* < 0.01.

| Overlap | Site | Station | Coy rel. abun. | Raccoon cluster | Coyote cluster |
|---|---|---|---|---|---|
| Ovr1 | DMP | RE 60 | LOW | | |
| Ovr1 | DMP | RW 100 | HIGH | | |
| Ovr1 | DMP | RW 117 | LOW | | |
| Ovr1 | DMP | RW 121 | HIGH | | |
| Ovr1 | DMP | RW 126 | HIGH | | |
| Ovr1 | DMP | RW 50 | HIGH | | |
| Ovr1 | DMP | RWC 1 | LOW | | |
| Ovr1 | DMP | RWC 6 | HIGH | | |
| Ovr1 | HMC | H17RE102 | LOW | | |
| Ovr1 | HMC | H17RW34 | LOW | | |
| Ovr1 | HMC | H17RW46 | HIGH | | |
| Ovr1 | HMC | H17RW47NEW | HIGH | | |
| Ovr2 | HMC | H17RE102 | LOW | | |
| Ovr2 | HMC | H17RE109 | HIGH | | |
| Ovr2 | HMC | H17RE123 | HIGH | | |
| Ovr2 | HMC | H17RW37 | HIGH | | |
| Ovr2 | HMC | H17RW43 | LOW | | |
| Ovr2 | HMC | H17RW47NEW | HIGH | | |
| Ovr2 | SNWR | RE 10 | LOW | | |
| Ovr2 | SNWR | RE 11 | LOW | | |
| Ovr2 | SNWR | RE 18 | HIGH | | |
| Ovr2 | SNWR | RE 8 | LOW | | |
| Ovr2 | SNWR | RW 10 | LOW | | |
| Ovr2 | SNWR | RW 11 | LOW | | |

Hot spot ($p < 0.01$)
Hot spot ($p < 0.05$)
Hot spot ($p < 0.1$)
Non-significant cluster
Cold spot ($p < 0.1$)
Cold spot ($p < 0.05$)
Cold spot ($p < 0.01$)

*Manuscript currently in revision at Ecosphere (as of June 2020)*

| | | | |
|---|---|---|---|
| Ovr2 | SNWR | RW 12 | LOW |
| Ovr2 | SNWR | RW 17 | HIGH |
| Ovr2 | SNWR | RW 24 | HIGH |
| Ovr2 | SNWR | RW 9 | LOW |
| Ovr2 | SNWR | RWC 1 | HIGH |
| Ovr2 | SNWR | RWC 2 | LOW |
| Ovr2 | SNWR | SWAP 100 | LOW |
| Ovr3 | HMC | H17RE106 | LOW |
| Ovr3 | HMC | H17RE127 | LOW |
| Ovr3 | HMC | H17RW34 | LOW |
| Ovr3 | HMC | H17RW37 | HIGH |
| Ovr3 | HMC | H17RW39 | LOW |
| Ovr3 | HMC | H17RW42 | LOW |
| Ovr3 | SNWR | RE 111 | LOW |
| Ovr3 | SNWR | RE 115 | HIGH |
| Ovr3 | SNWR | RE 118 | LOW |
| Ovr3 | SNWR | RE 120 | HIGH |
| Ovr3 | SNWR | RE 126 | LOW |
| Ovr3 | SNWR | RE 26 | LOW |
| Ovr3 | SNWR | RW 01 | LOW |
| Ovr3 | SNWR | RW 104 | HIGH |
| Ovr3 | SNWR | RW 113 | LOW |
| Ovr3 | SNWR | RW 117 | HIGH |
| Ovr3 | SNWR | RW 122 | LOW |
| Ovr3 | SNWR | RWC 08 | LOW |
| Ovr4 | SNWR | RE 111 | LOW |
| Ovr4 | SNWR | RE 118 | LOW |
| Ovr4 | SNWR | RE 120 | HIGH |
| Ovr4 | SNWR | RE 126 | LOW |
| Ovr4 | SNWR | RE 26 | LOW |

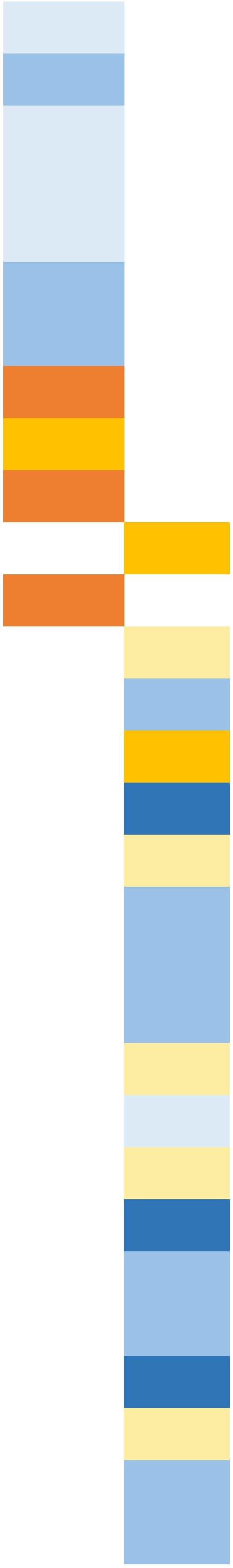

| | | | |
|---|---|---|---|
| Ovr4 | SNWR | RW 01 | LOW |
| Ovr4 | SNWR | RW 113 | LOW |
| Ovr4 | SNWR | RW 122 | LOW |
| Ovr4 | SNWR | RWC 08 | LOW |
| Ovr4 | UMBS | URE 12 | HIGH |
| Ovr4 | UMBS | URE 13 | HIGH |
| Ovr4 | UMBS | URE 2 | LOW |
| Ovr4 | UMBS | URW 1 | LOW |
| Ovr4 | UMBS | URW 23 | HIGH |
| Ovr4 | UMBS | URW 29 | LOW |
| Ovr4 | UMBS | URW 35 | LOW |
| Ovr4 | UMBS | URW 9 | HIGH |
| Ovr4 | UMBS | URWC 6 | HIGH |
| Ovr5 | SNWR | RE 26 | LOW |
| Ovr5 | SNWR | RW 01 | LOW |
| Ovr5 | SNWR | RW 112 | LOW |
| Ovr5 | UMBS | RE 10 | LOW |
| Ovr5 | UMBS | RE 15 | LOW |
| Ovr5 | UMBS | RE 16 | LOW |
| Ovr5 | UMBS | RE 18 | HIGH |
| Ovr5 | UMBS | RE 23 | HIGH |
| Ovr5 | UMBS | RE 24 | HIGH |
| Ovr5 | UMBS | RE 3 | HIGH |
| Ovr5 | UMBS | RE 4 | LOW |
| Ovr5 | UMBS | RE 5 | LOW |
| Ovr5 | UMBS | RW 10 | LOW |
| Ovr5 | UMBS | RW 11 | LOW |
| Ovr5 | UMBS | RW 15 | LOW |
| Ovr5 | UMBS | RW 23 | LOW |
| Ovr5 | UMBS | RW 4 | LOW |

| | | | |
|---|---|---|---|
| Ovr5 | UMBS | RW 9 | HIGH |
| Ovr5 | UMBS | RW16 | LOW |
| Ovr5 | UMBS | SWAP 100 | HIGH |
| Ovr6 | HMC | H17RE106 | LOW |
| Ovr6 | HMC | H17RE118 | HIGH |
| Ovr6 | HMC | H17RE127 | LOW |
| Ovr6 | UMBS | RE 1 | LOW |
| Ovr6 | UMBS | RE 10 | LOW |
| Ovr6 | UMBS | RE 15 | LOW |
| Ovr6 | UMBS | RE 23 | HIGH |
| Ovr6 | UMBS | RE 24 | HIGH |
| Ovr6 | UMBS | RE 5 | LOW |
| Ovr6 | UMBS | RE 8 | HIGH |
| Ovr6 | UMBS | RW 11 | LOW |
| Ovr6 | UMBS | RW 12 | LOW |
| Ovr6 | UMBS | RW 2 | HIGH |
| Ovr6 | UMBS | RW 22 | LOW |
| Ovr6 | UMBS | RW 23 | HIGH |
| Ovr6 | UMBS | RW 4 | LOW |
| Ovr6 | UMBS | RW 8 | LOW |
| Ovr6 | UMBS | RW 9 | HIGH |
| Ovr6 | UMBS | RW16 | LOW |
| Ovr7 | HMC | H17RE127 | LOW |
| Ovr7 | HMC | H17RE18 | LOW |
| Ovr7 | HMC | H17RW34 | LOW |
| Ovr7 | HMC | H17RW37 | HIGH |
| Ovr7 | HMC | H17RW39 | LOW |
| Ovr7 | HMC | H17RW42 | HIGH |
| Ovr7 | UMBS | URE 12 | HIGH |
| Ovr7 | UMBS | URE 2 | LOW |

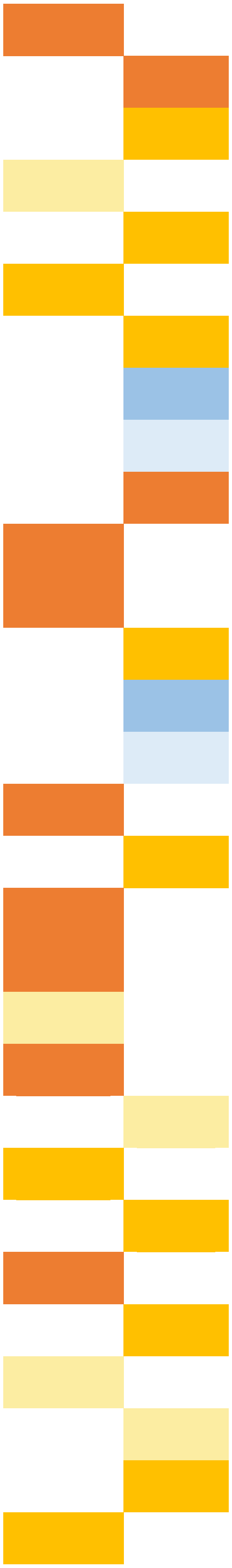

*Manuscript currently in revision at Ecosphere (as of June 2020)*

| | | | |
|---|---|---|---|
| Ovr7 | UMBS | URE 27 | LOW |
| Ovr7 | UMBS | URW 1 | LOW |
| Ovr7 | UMBS | URW 15 | HIGH |
| Ovr7 | UMBS | URW 23 | HIGH |
| Ovr7 | UMBS | URW 29 | LOW |
| Ovr7 | UMBS | URW 35 | LOW |
| Ovr7 | UMBS | URW 5 | HIGH |
| Ovr7 | UMBS | URWC 6 | HIGH |
| Ovr8 | HMC | RE 24 NEW | HIGH |
| Ovr8 | HMC | RE 25 | HIGH |
| Ovr8 | HMC | RE 39 | LOW |
| Ovr8 | HMC | RE 4 | LOW |
| Ovr8 | HMC | RE 48 | HIGH |
| Ovr8 | HMC | RE 50 | HIGH |
| Ovr8 | HMC | RE 6 | LOW |
| Ovr8 | HMC | RW 21 | HIGH |
| Ovr8 | HMC | RW 22 | HIGH |
| Ovr8 | HMC | RW 24 | HIGH |
| Ovr8 | HMC | RW 3 | LOW |
| Ovr8 | HMC | RW 36 | LOW |
| Ovr8 | UMBS | URE 2 | LOW |
| Ovr8 | UMBS | URW 15 | HIGH |
| Ovr8 | UMBS | URW 23 | HIGH |
| Ovr8 | UMBS | URW 29 | LOW |
| Ovr8 | UMBS | URW 35 | LOW |
| Ovr8 | UMBS | URW 9 | HIGH |
| Ovr8 | UMBS | URWC 4 | HIGH |
| Ovr8 | UMBS | URWC 6 | HIGH |
